# Spatial and temporal profiling of receptor membrane insertion controls commissural axon responses to midline repellents

**DOI:** 10.1101/540690

**Authors:** Aurora Pignata, Hugo Ducuing, Leila Boubakar, Thibault Gardette, Karine Kindbeiter, Muriel Bozon, Servane Tauszig-Delamasure, Julien Falk, Olivier Thoumine, Valérie Castellani

**Affiliations:** University of Lyon, University of Lyon 1 Claude Bernard Lyon1, NeuroMyoGene Institute, CNRS UMR5310, INSERM U1217, 8 Avenue Rockefeller, 69008 Lyon, France; Interdisciplinary Institute for Neuroscience, UMR CNRS 5297 - University of Bordeaux 146 rue Léo Saignat, F-33000 Bordeaux, France

## Abstract

Accurate perception of guidance cues is crucial for axonal pathfinding. During their initial navigation in the spinal cord, commissural axons are kept insensitive to midline repellents. Through yet unclear mechanisms acting during midline crossing in the floor plate, they switch on responsiveness to various repulsive signals, that establish a permanent midline barrier and propel the axons for exit. Whether these gains of response are coupled to occur in synchrony or rather are independently activated through signaling-specific programs is fully unknown. We set-up a paradigm for live imaging and super resolution analysis of guidance receptor dynamics during commissural growth cone navigation in chick and mouse embryos. We uncovered a remarkable program of delivery and allocation of receptors at the growth cone surface, generating receptor-specific spatial and temporal profiles. This reveals a mechanism whereby commissural growth cones can discriminate coincident repulsive signals that they functionalize at different time points of their navigation.

## Introduction

In a broad range of biological contexts, cells are exposed to a complex array of environmental cues from which they receive specific instructions. This is well exemplified by the model of axon responses to guidance cues during the formation of neuronal circuits. Axons navigate highly diverse environments to reach their targets. Unique trajectories emerge from the perception by axon tips, the growth cones, of combinations of extracellular cues exposed in choice points along the paths. A typical case is provided by commissural neurons, which must project their axons across the midline to build with contralateral target cells circuits integrating left and right neuronal activities (Evans and Bashaw, 2010; Pignata *et al*., 2016; Stoeckli, 2018). Midline crossing of commissural axons in the floor plate (FP) of the developing spinal cord has been extensively worked out to explore axon guidance mechanisms, especially those regulating growth cone sensitivity to guidance cues. Various repulsive forces provided by proteins of the Slit and Semaphorin families are needed to prevent midline re-crossing and expel the axons away towards their next step. They are also thought to control the lateral position relative to the FP that commissural axons take to navigate their rostrally-oriented longitudinal path after the crossing (Long *et al*., 2004; Jaworski *et al*., 2010). Semaphorin3B (Sema3B) acting via Neuropilin2 (Nrp2)-PlxnA1 receptor complex, N-terminal and C-terminal Slit fragments resulting from Slit processing acting respectively via Roundabout (Robo)1/2 and PlexinA1 (PlxnA1) receptors are guidance cues all found expressed in the FP and playing instructive roles during commissural axon navigation (Zou *et al.*, 2000; Long *et al.*, 2004; Jaworski *et al.*, 2010; Nawabi *et al.*, 2010; Delloye-Bourgeois *et al.*, 2015).

Manipulations of Semaphorin and Slit repulsive signaling in mouse and chicken embryo models brought the view that the sensitivity of commissural axons to midline repellents must be silenced in a first step, prior to the crossing, and switched on in a second step to allow repulsive forces to set a midline barrier and expel the growth cones away. Consistently, manipulations presumably inducing premature sensitization or preventing it resulted in failure of FP crossing, with axons arrested before or within the FP, turning back or longitudinally before reaching the contralateral side (Chen *et al.*, 2008; Nawabi *et al.*, 2010). Noteworthy, the FP navigation is not a synchronous process. For example, it extends over several days in the mouse embryo, from the first axon wave of earliest-born commissural neurons at E9.5 to the latest one at E12.5 (Wilson *et al*., 2008; Pignata *et al*., 2016). The repellents are expressed in the FP over the entire period of commissural navigation (Pignata *et al.*, 2016). Thus, independently from the ligand expression profiles, the switch towards sensitivity upon crossing has to be set at individual growth cone level. Whether the sensitization of commissural axons to the various repellents they encounter in the FP occurs in synchrony, or rather presents signaling-specific features is unknown. Also, whether repulsive guidance receptors distribute homogeneously or present spatial specificities at the growth cone surface is undetermined. Insights are still scarce essentially due to the deficit of experimental paradigms giving access to molecular events in single living commissural axons navigating in their native context.

We postulated that functional engagement of midline repellents could arise from peculiar dynamics of guidance receptors at the surface of navigating growth cones. To address this hypothesis, we investigated the cell surface dynamics of four receptors mediating repulsion by midline cues, Nrp2, PlxnA1, Robo1 and Robo2, in chick and mouse embryo models. Surprisingly, we found striking differences between Robo1 and Robo2 temporal patterns, which excludes Robo2 as a mediator of Slit repulsion during FP crossing but places it as a major player of the lateral funiculus navigation. Our study also revealed exquisite specificities of PlxnA1 and Robo1 dynamics. Both receptors are not only sorted at different timing of FP navigation but also are distributed in distinct domains of the growth cones. This spatial and temporal compartmentalization is achieved at post-translational and post-intra-axonal trafficking levels, specifically at the step of membrane delivery in the growth cones. Analysis of the dynamics of PlxnA1-Robo1 chimeric receptors demonstrated that the intracellular domain of PlxnA1 but not that of Robo1, is sufficient for coding the receptor-specific temporal pattern. Finally, FRAP analysis in growth cones navigating the FP further confirmed dynamics specificities of these two receptors. Our study thus revealed remarkably unique spatial and temporal sequences of cell surface sorting of guidance receptors during the navigation of spinal commissural axons. This mechanism enables the growth cones to discriminate in time and space coincident guidance signals and provides a basis for these cues to exert non-redundant and concerted actions.

## Results

### Development of an experimental paradigm to visualize cell surface receptor dynamics in navigating commissural axons

We setup time lapse imaging to monitor the cell surface dynamics of Semaphorin and Slit receptors in commissural axons navigating the FP in native spinal cords of chicken embryos. Nrp2, PlxnA1, Robo1 and Robo2 receptors were fused to the pH-sensitive GFP, pHLuorin (pHLuo), whose fluorescence at neutral pH enables to report membrane protein pools and cloned in vectors with ires-mb-tomato (**Fig. 1a**) (Jacob *et al.,* 2005; Nawabi *et al.*, 2010; Delloye-Bourgeois *et al.*, 2014). The pH-dependency of receptor fluorescence was verified by *in vitro* cell-line transfections (**Supplementary Fig. 1a**) (Delloye-Bourgeois *et al.*, 2014). The vectors were then transferred to spinal cord commissural neurons using *in ovo* neural tube electroporation (**Fig. 1b**). Then, isolated spinal cords were dorsally opened and imaged over several hours for mapping the receptor cell surface sorting reported by pHLuo fluorescence. The FP entry and exit limits were delineated using DIC channel or based on the observation of some typical features of axon trajectory, such as the presence of wrinkles when axons enter the FP and the rostral turning when axons exit the FP (**Supplementary Fig. 1b**).

**Fig. 1:**
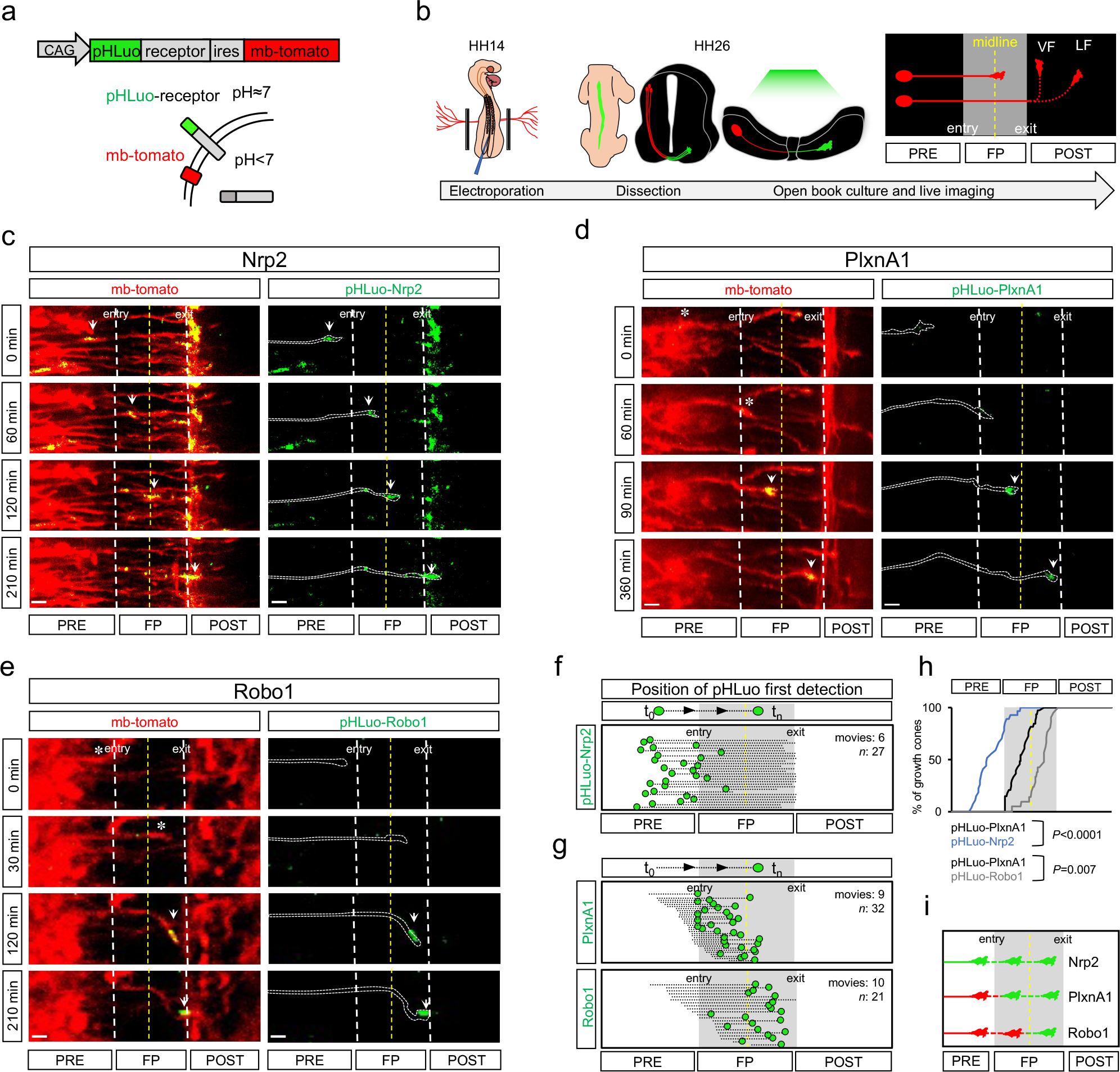
PlxnA1 and Robo1 are successively sorted to the growth cone surface during FP navigation. **a**) Schematic drawings of pHLuo vector and pH-dependent pHLuo fluorescence activity in cells reporting cell surface protein pool. pHLuorin-receptor and mb-tomato coding sequences were cloned in pCAG vector, spaced by an ires. (**b**) *In ovo* electroporation procedure. 48h after electroporation, spinal cords were dissected and mounted as open books for time lapse microscopy. pHLuo fluorescence was monitored in 3 compartments of the open-books: the pre-crossing, where the cell bodies of commissural neurons are located, the FP, in which commissural axons cross the midline, and the post-crossing, in which commissural axons chose between ventral and lateral longitudinal trajectories. VF: ventral funiculus, LF: lateral funiculus. (**c-e**) Time-lapse sequences illustrating the spatial and temporal dynamics of pHLuo-Nrp2, pHLuo-PlxnA1 and pHLuo-Robo1 during FP navigation. The asterisks point growth cone positions before pHLuo flashes and the white arrowheads those of pHLuo flashes and subsequent growth cone positions. (**f**) Cartography of pHLuo-Nrp2 dynamics from movie analysis. Dashed lines indicate the overall trajectory of single growth cones and green spots the first pHLuo detection. Nrp2 is exposed at the growth cone surface since the onset of spinal cord navigation (Nrp2: N=5 embryos, 6 movies, 27 growth cones). (**g**) Cartography of pHLuo-PlxnA1 and pHLuo-Robo1 dynamics, plotting position of pHLuo flashes. The upper panel illustrates pHLuo-PlxnA1 sorting in the first FP half, the lower panel that of pHLuo-Robo1 in the second FP half (PlxnA1: N=5 embryos, 9 movies, 32 growth cones; Robo1: N= 9 embryos, 10 movies, 21 growth cones). (**h**) Cumulative fractions showing differential pHLuo-Nrp2, pHLuo-PlxnA1 and pHLuo-Robo1 dynamics during FP navigation (*P* value is from the Kolmogorov-Smirnov (KS) test). (**i**) Summary of the temporal sequence of pHLuo-Nrp2, pHLuo-PlxnA1 and pHLuo-Robo1 membrane sorting during FP navigation. Scale bars in **c-e**, 10 μm.

### PlxnA1 and Robo1 are specifically and successively sorted to the growth cone surface during FP navigation

Using our setup, we analyzed individual growth cone trajectories from time-lapse sequences. We plotted the position of the growth cones that turned on the pHLuo fluorescence to build cartographies of receptor cell surface sorting positions along the navigation. First, we observed that Nrp2 is exposed at the commissural growth cone surface from the pre-crossing stage and remains over entire FP crossing (**Fig. 1c,f,h,i; Supplementary Movie 1,2**). In contrast, we found that the membrane sorting of both PlxnA1 and Robo1 specifically occurs during FP navigation. Interestingly, the timing of their sorting significantly differed. PlxnA1 was addressed to the surface when commissural growth cones navigate the first half of the FP, thus from the FP entry point to the midline (**Fig. 1d,g,h,i; Supplementary Movie 3,4**), while Robo1 was sorted during the navigation of the second FP half, from the midline to the FP exit point (**Fig. 1e,g,h,i; Supplementary Movie 5,6**).

### PlxnA1 and Robo1 temporal patterns are set by control of cell surface sorting

Next, we assessed whether these temporal patterns are profiled by control of receptor cell surface sorting or rather by control of protein availability within the axon. Spinal cord open books were fixed with paraformaldehyde (PFA) at neutral pH to detect the total pHLuo receptor pool. While in live axons pHLuo signal was found restricted to the growth cones during FP crossing, we observed in contrast for both receptors that the total pools had much broader distribution than the surface ones. In 90% of the cases for PlxnA1 and 75% for Robo1, the pre-crossing axon segment immediately adjacent to the FP entry of growth cones that were found to navigate within the FP, contained pHLuo-receptors (**Fig. 2a,b**). We also measured pHLuo^+^ pre-crossing segment length in the fixed samples and found significant difference between PlxnA1 and Robo1, the latter having more restricted length and punctate pattern than the first one (**Fig. 2c**). These observations are consistent with previous works that reported in cultured commissural neurons Robo1 intra-axonal vesicular trafficking (Philipp *et al.*, 2012) and PlxnA1 processed within axons to prevent its membrane expression (Nawabi *et al.*, 2010). Thus, PlxnA1 and Robo1 are available within commissural axons and their cell surface sorting is spatially and temporally controlled in a receptor-specific manner.

**Fig. 2:**
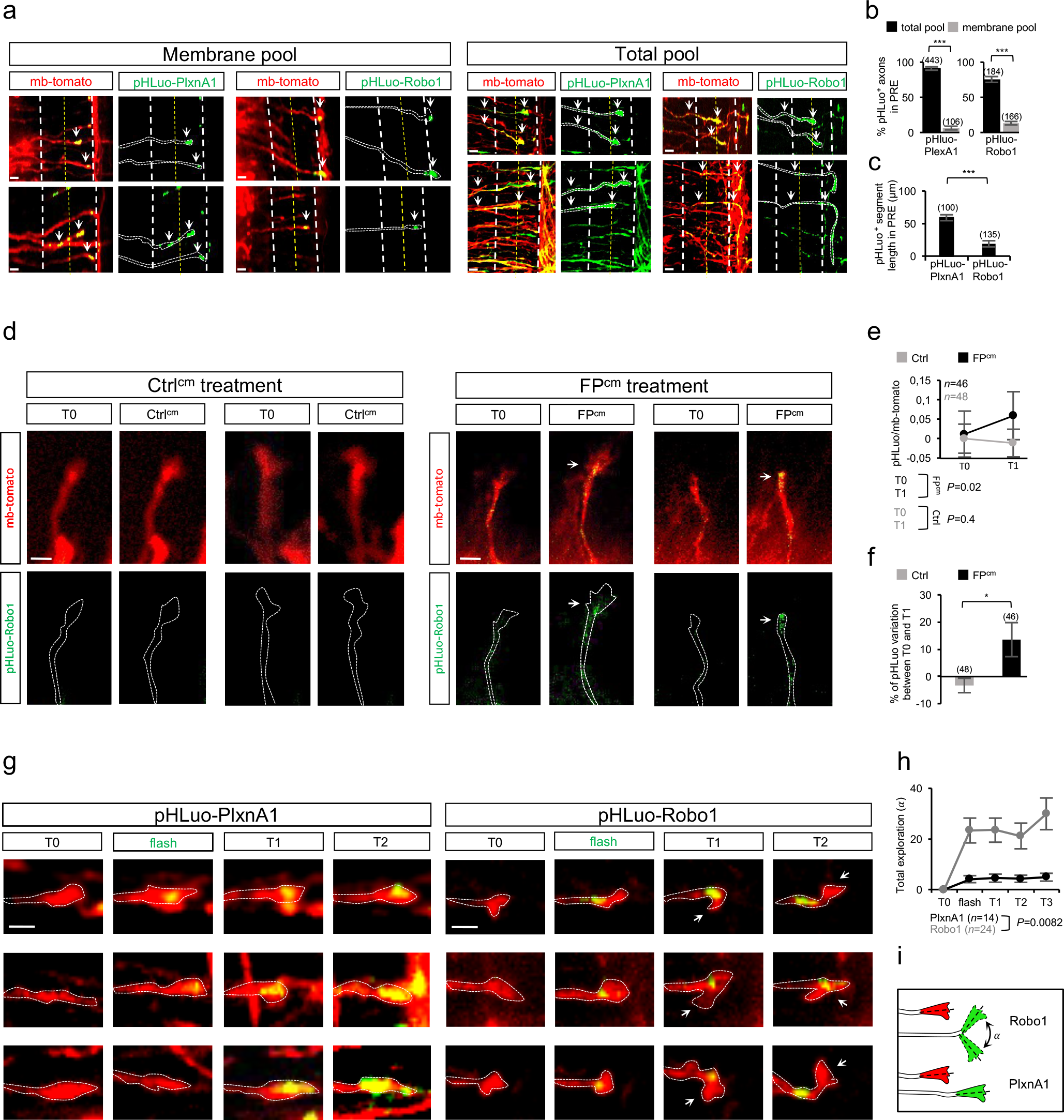
Cell surface sorting of PlxnA1 and Robo1 is temporally controlled and coincident with changes of exploratory behavior for Robo1. **a**) Microphotographs of open books illustrating pHLuo-PlxnA1 and pHLuo-Robo1 membrane pool (left panel) and total (intracellular + membrane) receptor pool (right panel). Arrowheads point discrete pHLuo^+^ growth cones and axon segments. (**b**) Quantification of the % of growth cone population observed to navigate within the FP and containing pHLuo-receptor in the pre-crossing segment immediately adjacent to the FP entry. Histograms show much broader total than surface fluorescence in this segment (total PlxnA1: N=4 embryos, 443 growth cones; membrane PlxnA1: N=5 embryos, 106 growth cones; total Robo1: N=4 embryos, 184 growth cones; membrane Robo1: N=17 embryos, 166 growth cones. Error bars indicate mean ± SEM; p <0.05; ** p <0.01; *** p <0.001. *P* values are from Mann-Whitney test). (**c**) Histograms of measured lengths of pre-crossing segment containing total pHLuo-PlxnA1 or pHLuo-Robo1 in growth cones observed to navigate the FP, showing a more restricted pHLuo-Robo1 pattern, compared to that of pHLuo-PlxnA1. (**d**) Electroporated dorsal explant cultures treated with Ctrl^cm^ (left panel) or FP^cm^ (right panel) showing pHLuo-Robo1 increase at the growth cone membrane after FP^cm^ application. (**e**) Quantitative analysis showing the increase of pHLuo-Robo1 at the growth cone surface after 20 min (T1) of FP^cm^ treatment. For each growth cone, pHLuo is normalized to mb-tomato signal (3 independent experiments; Ctrl: N=19 explants, 48 growth cones; FP^cm^: N=18 explants, 46 growth cones. Error bars indicate mean ± SEM; * p <0.05; ** p <0.01; *** p <0.001. *P* values are from paired Student t test). (**f**) Quantification of pHLuo-Robo1 signal variation between T0 and T1 in Ctrl^cm^ and FP^cm^ conditions, showing the increase of surface Robo1 after FP^cm^ application (Error bars indicate mean ± SEM; * p <0.05; ** p <0.01; *** p <0.001. *P* values are from unpaired Student t test). (**g**) Photomicrographs of pHLuo-PlxnA1 (left panel) and pHLuo-Robo1 (right panel) sorting at the growth cone membrane. Arrowheads in pHLuo-Robo1 condition point the exploratory behavior of growth cones after pHLuo sorting. (**h**) Quantitative analysis of the total angle explored by growth cones from the time point just preceding the flash (T0) to 1,5 hours after the flash (T3) (PlxnA1: N= 3 embryos, 32 growth cones; Robo1: N=10 embryos, 21 growth cones. Error bars indicate mean ± SEM. P value is from the Kolmogorov-Smirnov (KS) test). (**i**) Graphic summary of the exploratory behavior of the growth cone following pHluo-Robo1 surface sorting. Scale bars in **a** and **d**, 10 μm; scale bars in **g**, 5 μm.

We found in previous work that medium conditioned by cultured isolated FP tissues (FP^cm^) could trigger PlxnA1 cell surface expression (Nawabi *et al.*, 2010). Such medium was also reported to induce in commissural growth cones Robo3.2, the Robo3 isoform expressed in post-crossing axons (Colak *et al*., 2013), providing the evidence that local signals emanating from the FP are implicated in synchronizing the sorting of these receptors with midline crossing. How is triggered the sorting of Robo1 is yet unknown. We thus examined whether it could also be under local FP control. We treated dorsal spinal cord explants electroporated with pHLuo-Robo1-ires-mb-tomato with FP^cm^ and ctrl medium and recorded Robo1 dynamics by measuring pHLuo fluorescence in the growth cones at T0 and T1, 20 min later (**Fig. 2d**). We observed a significant increase of pHLuo fluorescence at T1 compared to T0 for the FP^cm^ but not the control condition (**Fig. 2e,f**), thus indicating that FP cells release cues triggering Robo1 at the growth cone surface.

Next, we assessed whether disturbing the temporal pattern of receptor sorting impacts on growth cone behaviors. Open-books were electroporated with high concentration of vectors (3μg/μl and 4μg/μl) to overcome the internal control of PlxnA1 and Robo1 surface sorting in commissural neurons and create premature surface expressions. We monitored individual growth cones and found that commissural growth cones having premature cell surface receptor exposure failed to cross the FP, rather turning or stalling before or within the FP (**Supplementary Fig. 2; Supplementary Movie 7,8**). These findings indicated that pHLuo-receptors are functional and confirmed that the temporal pattern of receptor sorting is critical for proper FP navigation.

Next, we asked whether the gain of PlxnA1 and Robo1 at the surface could be correlated with acquisition of novel behavioral properties of the navigating growth cones. To address this question, we analyzed in time-lapse movies growth cone trajectories at time-points preceding and succeeding the pHLuo flashes, by measuring the deviation angles of growth cone direction from the trajectory baseline (**Fig. 2g**). Interestingly for Robo1, we observed that acquisition of surface receptor was coincident with a significant increase of exploratory behavior along the rostro-caudal axis as if the growth cones were starting sensing cues that will direct their longitudinal turning at the FP exit. In contrast, we found no difference of exploration for PlxnA1 (**Fig. 2g,h,i**).

### Robo2 is sorted not in the FP but in the post-crossing lateral funiculus

Next, we investigated the dynamics of Robo2. In sharp contrast with Robo1, we found that Robo2 was absent from the surface of commissural growth cones navigating the FP and turning longitudinally at medial position in the ventral funiculus (VF). Instead, we observed that it was specifically sorted in post-crossing axons that chose to navigate in the lateral funiculus and turned longitudinally to the FP (LF) (**Fig. 3a,b,c; Supplementary Movie 9,10**). To assess if Robo2 cell surface sorting correlates with this change of trajectory, we measured the angle formed by a vector aligned along the axon tip and the FP axis at the two time points framing Robo2 pHLuo flash. Interestingly, we found that the angle was significantly more pronounced at post-flash time points compared with pre-flash ones, supporting that Robo2 sorting contributes to directional growth cone changes along the longitudinal axis (**Fig. 3d,e**). Thus interestingly, signaling by Robo1 and Robo2 appear to have similar outcome on growth cone behaviors, that they control at two different time points of commissural navigation.

**Fig. 3:**
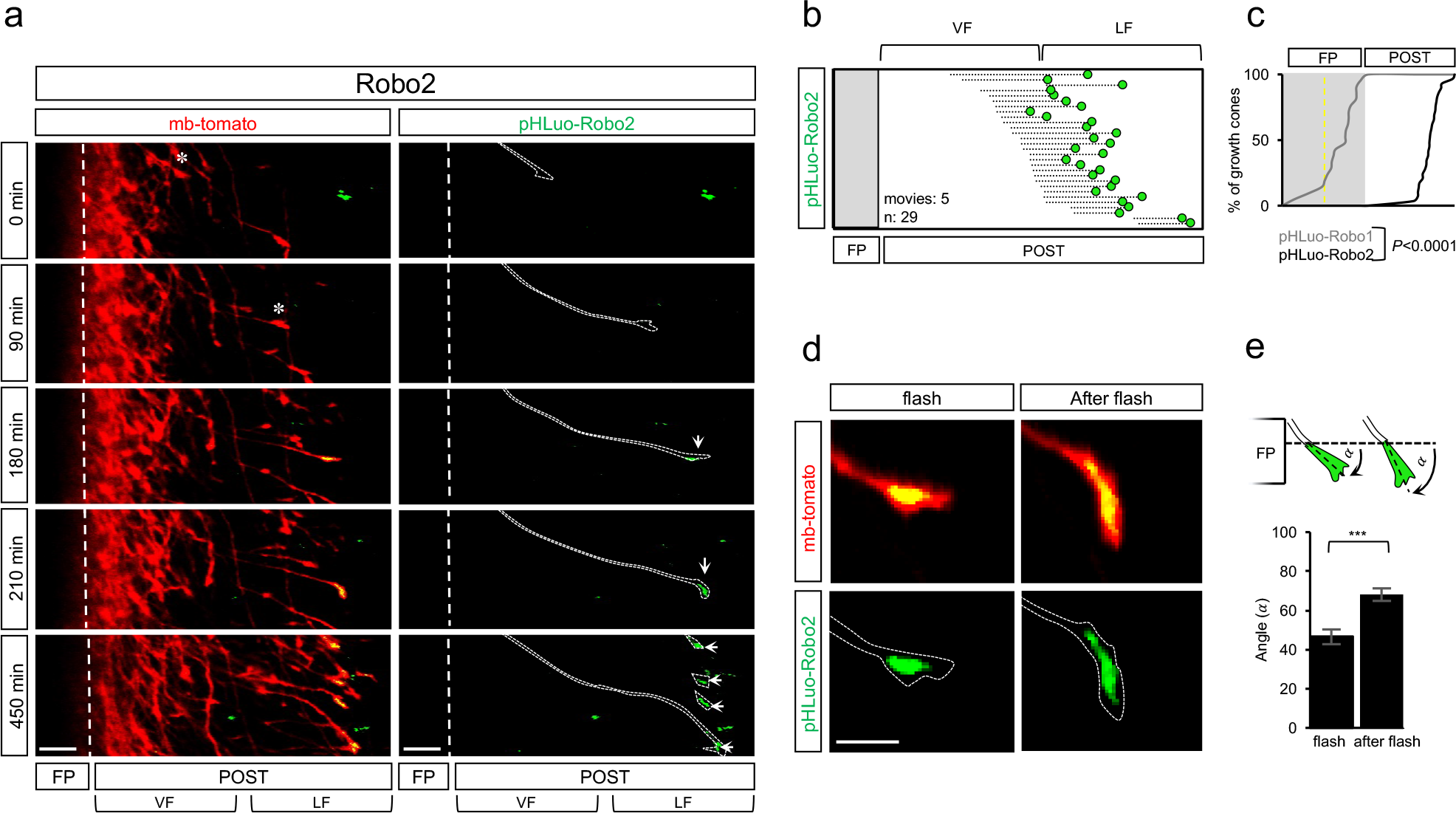
Robo2 is not sorted in growth cones navigating the FP but during post-crossing pathfinding of the lateral funiculus. (**a**) Time-lapse sequences of open-books illustrating pHLuo-Robo2 spatio-temporal dynamics in commissural growth cones. The asterisks point growth cone positions before pHLuo flashes and the white arrowheads those of pHLuo flashes and subsequent growth cone positions. (**b**) Cartography of pHLuo-Robo2 flashes. Dashed lines indicate the overall trajectory of individual growth cones, from imaging onset to the time point of flash occurrence (Robo2: N=4 embryos, 5 movies, 29 growth cones). (**c**) Cumulative fractions showing differential pHLuo-Robo1 and pHLuo-Robo2 dynamics during spinal cord navigation (*P* value is from the Kolmogorov-Smirnov (KS) test). (**d**) Representative images from time-lapse sequence illustrating a shift of growth cone orientation subsequent to pHLuo-Robo2 flash. (**e**) Schematic drawing and quantification of growth cone turning after pHLuo-Robo2 flashes (Robo2: N=3 embryos, 30 growth cones. Error bars indicate mean ± SEM; * p <0.05; ** p <0.01; *** p <0.001. *P* value is from Mann-Whitney test). Scale bars in **a**, 50 μm, in **d**, 10 μm.

### The temporal sequence of receptor sorting is conserved in the mouse

Next, we studied whether the temporal control of receptor sorting during FP navigation uncovered in the chicken embryo is conserved in the mouse and whether it also instructs growth cone guidance choices. We electroporated pHLuo-PlxnA1 and pHLuo-Robo1 constructs in the developing spinal cord of E12 wild-type mouse embryos. We plotted the position of fluorescent growth cones in living open-books at fixed time point, 48 hours post-electroporation, when many FP crossing are ongoing, as depicted by the distribution of mb-tomato^+^ growth cones (**Fig. 4c,d**). In pHLuo-PlxnA1 electroporated spinal cords, growth cones exposing the pHLuo were distributed almost homogenously in all FP and post-crossing compartments (**Fig. 4a,b,c**) whereas in the pHLuo-Robo1 electroporated littermates, most of the growth cones exposing Robo1 were situated between the midline and FP exit (**Fig4. a,d,e**). Therefore, the spatial and temporal cell surface pattern of PlxnA1 and Robo1 observed in chick spinal cord is conserved in mice (**Fig. 4e**).

**Fig. 4:**
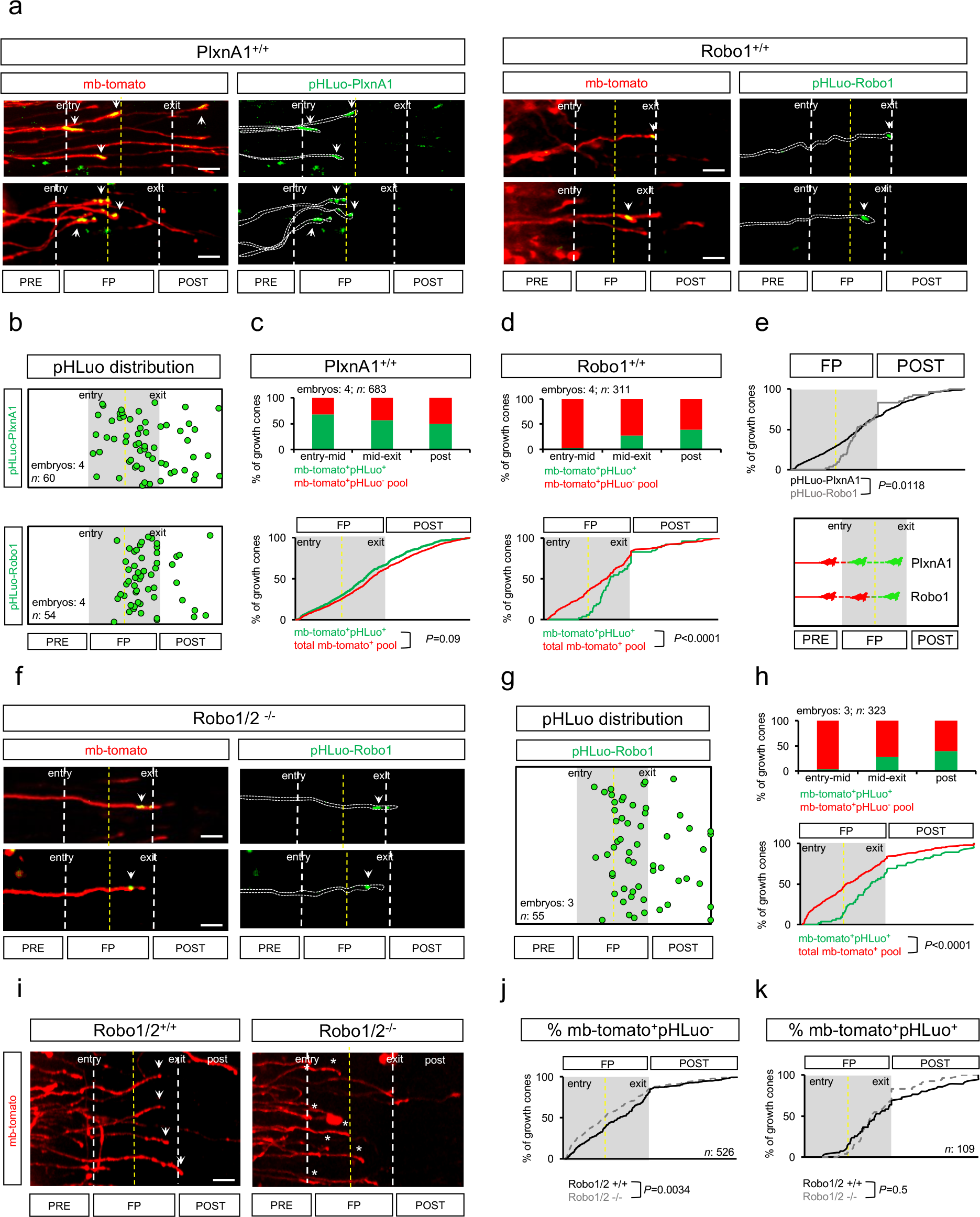
chick PlxnA1 and Robo1 temporal sequences are conserved in the mouse. Microphotographs of PlxnA1^+/+^ and Robo1^+/+^ open-books illustrating pHLuo-PlxnA1^+^ (left panel) and pHLuo-Robo1^+^ (right panel) growth cones pointed by white arrowheads. (**b**) Upper panel: cartography of pHLuo-PlxnA1^+^ growth cones (N=4 embryos, 60 growth cones); lower panel: cartography of pHLuo-Robo1^+^ growth cones (N=4 embryos, 54 growth cones). (**c**) Upper panel: distribution of mb-tomato^+^pHLuo^+^ and mb-tomato^+^pHLuo^−^ populations in PlxnA1^+/+^ open books. Lower panel: cumulative fraction of mb-tomato^+^pHLuo^+^ and total mb-tomato^+^ (composed of mb-tomato^+^pHLuo^+^ and mb-tomato^+^pHLuo^−^) populations showing that mb-tomato^+^pHLuo^+^ growth cones are detected from the onset of FP navigation (*P* value is from the Kolmogorov-Smirnov (KS) test). (**d**) Upper panel: distribution of mb-tomato^+^pHLuo^+^ and mb-tomato^+^pHLuo^−^ populations in the spinal cord of Robo1/2^+/+^ open books. Lower panel: cumulative fractions of mb-tomato^+^pHLuo^+^ and total mb-tomato^+^ (composed of mb-tomato^+^pHLuo^+^ and mb-tomato^+^pHLuo^−^) populations showing that whereas the total mb-tomato^+^ population distributes from the first FP half to the post-crossing compartment, mb-tomato^+^pHLuo^+^ growth cones are only detected from the second half of the FP (*P* value is from the Kolmogorov-Smirnov (KS) test). (**e**) Cumulative fraction of pHLuo-PlxnA1^+^ and pHLuo-Robo1^+^ growth cones, showing the differential timing of sorting of the receptors in the FP (*P* value is from the Kolmogorov-Smirnov (KS) test). (**f**) Microphotographs of Robo1/2^−/−^ open-books illustrating pHLuo-Robo1^+^ growth cones pointed by white arrowheads. (**g**) Cartography of pHLuo-Robo1^+^ growth cones (N=3 embryos, 55 growth cones). (**h**) Upper panel: distribution of mb-tomato^+^pHLuo^+^ and the mb-tomato^+^pHLuo^−^ populations in Robo1/2^−/−^ open books. Lower panel: cumulative fractions of mb-tomato^+^pHLuo^+^ and total mb-tomato^+^ (*P* value is from the Kolmogorov-Smirnov (KS) test). (**i**) Representative images of mb-tomato^+^ growth cones illustrating reduced number of growth cones (asterisks) on their way for FP exit in Robo1/2^−/−^ compared to Robo1/2^+/+^ open-books. (**j**) Cumulative fractions reporting the distribution of mb-tomato^+^pHLuo^−^ growth cones in Robo1/2^+/+^ and Robo1/2^−/−^ embryos, showing a significantly shifted distribution of tomato^+^pHLuo^−^ growth cones towards the first FP half in Robo1/2^−/−^ embryos. (**k**) Cumulative fractions reporting similar distribution of pHLuo-Robo1^+^ growth cones in open-books of Robo1/^+/+^ and Robo1/2^−/−^ embryos (*P* value is from the Kolmogorov-Smirnov (KS) test). Scale bars in **a** and **f** 20 μm, in **i**, 50 μm.

We then electroporated pHLuo-Robo1 construct in Robo1/2^−/−^open books to determine whether reducing Robo1 dose by removal of the endogenous pool results in a modified Robo1 temporal pattern. We found that the profile of receptor sorting was identical to that observed after electroporation of Robo1/2^+/+^ embryos (**Fig. 4f,g,h**). Thus, our experimental conditions of expression are likely to model the dynamics of endogenous receptor. This result also established that Robo1 sorting at the plasma membrane is Robo2 independent.

Next, we investigated whether the re-expression of pHLuo-Robo1 in Robo1/2^−/−^ mice could rescue the previously reported stalling phenotypes resulting from Robo1/2 deletion (Long *et al.*, 2004; Delloye-Bourgeois *et al.*, 2015). We analyzed the distribution of mb-tomato^+^ growth cones over pre-to post-crossing steps, distinguishing growth cones that exposed Robo1 at their surface (mb-tomato^+^pHLuo^+^) from those that did not (mb-tomato^+^pHLuo^−^) in Robo1/2^+/+^ and Robo1/2^−/−^ embryos. We observed that Robo1/2 loss resulted in significantly shifted distribution of mb-tomato^+^pHLuo^−^towards the first FP half (**Fig. 4i,j**). Interestingly, the expression of Robo1 at the growth cone surface was sufficient to rescue the distribution observed in the WT condition, as observed by the matching of the distribution of mb-tomato^+^pHLuo^+^ growth cones in Robo1/2^−/−^ and Robo1/2^+/+^ embryos (**Fig. 4k**). Thus, re-expression of Robo1 coding sequence in commissural neurons and subsequent cell surface exposure at a time when the growth cone navigates the second half of the FP is sufficient to rescue proper navigation. Moreover, and consistent with its observed temporal sorting profile, Robo2 is dispensable for FP crossing and Robo1 surface exposure. Thus, this supports that this temporal profile properly reports the dynamics of endogenous Robo1 receptor.

### PlxnA1 and Robo1 are partitioned at the growth cone surface

Next, we used super resolution microscopy to assess whether Robo1 and PlxnA1, on top of the different timing at which they are sorted during FP crossing, also differ in their spatial distribution at the growth cone surface. Living open-books were incubated with ATTO-647N-conjugated GFP nanobodies to label cell surface pHLuo. After fixation, receptor pools were imaged in commissural growth cones at different steps of their FP navigation, using STED microscopy (**Fig. 5a**). First, we measured the density of the fluorescent signal in individual growth cones. We found that PlxnA1 and Robo1 receptor clusters have differential cell surface distributions, PlxnA1 predominantly accumulating at the growth cone front, and Robo1 at the rear (**Fig. 5b,c,d**). This was also confirmed by determining the center of mass of the signals, which segregated along the growth cone front-rear axis (**Fig. 5e**). Second, we studied whether the distribution patterns of PlxnA1 and Robo1 vary over FP navigation. According to their temporal sorting, we compared PlxnA1 distributions between FP entry and exit, and those of Robo1 between midline and exit. Analysis of the number and the size of labelled individual particles revealed modifications of Robo1 but not PlxnA1 patterns (**Fig. 5f,g**). Although not differing in their numbers, the size of pHLuo-Robo1^+^ particles increased from the midline to the exit, indicative of Robo1 diffusion at the surface (**Fig. 5g**).

**Fig. 5:**
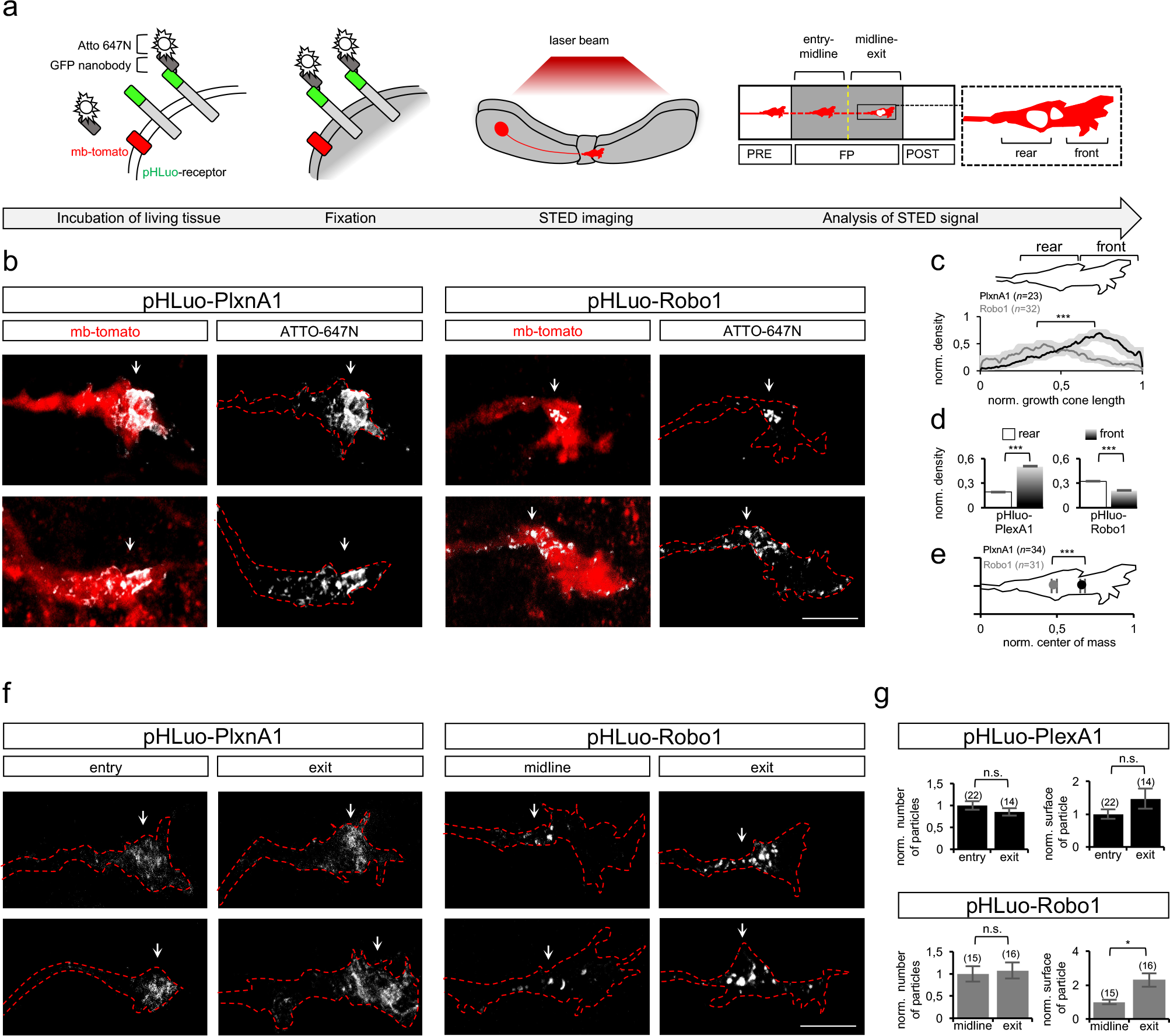
PlxnA1 and Robo1 are partitioned at the cell surface of commissural growth cones navigating the FP. Super resolution imaging procedure. Open-books of embryos electroporated with pHLuo-vectors were live-labelled using ATTO-647N-conjugated GFP nanobodies and were fixed with PFA before STED imaging. Membrane pHLuo density and distribution were analyzed in the growth cones navigating the first (entry-midline) and the second (midline-exit) halves of the FP. The growth cone was segmented into front and rear sub-domains. (**b**) Microphotographs of representative commissural growth cones delineated with mb-tomato and labeled with ATTO-647N-conjugated GFP nanobodies. White arrowheads point ATTO-647N signal. (**c**) Densities of membrane pHLuo-PlxnA1 and pHLuo-Robo1 signals normalized to the growth cone length, showing their differential distribution along the growth cone rear-front axis (PlxnA1: N=8 embryos, 23 growth cones; Robo1: N=12 embryos, 32 growth cones. Error bars indicate mean ± SEM; * p <0.05; ** p <0.01; *** p <0.001. *P* is from Kolmogorov-Smirnov (KS) test). (**d**) Histograms showing the comparison between normalized density of pHLuo signal in the front and the rear domain for both pHLuo-PlxnA1 and pHLuo-Robo1 (*P* value is from Mann-Whitney test). (**e**) Positions of pHLuo-PlxnA1 and -Robo1 center of mass normalized to growth cone lengths (PlxnA1: N=34 growth cones; Robo1: N=31 growth cones). (**f**) Microphotographs of mb-tomato^+^pHLuo^+^ commissural growth cones labeled with ATTO-647N-conjugated GFP nanobodies and navigating the entry-midline or the midline-exit compartments. (**g**) Upper panel: histograms showing normalized numbers and surfaces of pHLuo-PlxnA1 individual clusters detected in growth cones navigating the FP entry and exit. The number and the surface of pHLuo-PlxnA1 clusters were unchanged along FP navigation (PlxnA1: N=7 embryos, 22 growth cones in entry, 14 growth cones in exit (Error bars indicate mean ± SEM; * p <0.05; ** p <0.01; *** p <0.001. *P* value is from Mann-Whitney test). Lower panel: histograms showing the normalized numbers and surfaces of pHLuo-Robo1 individual clusters detected in growth cones navigating the FP midline and exit. Note that the particle number remained unchanged whereas the particle surface increased between midline and exit (Robo1: N=5 embryos, 15 growth cones in midline, 16 cones in exit. Error bars indicate mean ± SEM; * p <0.05; ** p <0.01; *** p <0.001. *P* value is from Mann-Whitney test). Scale bars in **b,f**, 5 μm.

We next investigated whether such compartmentalization is generated at the surface or rather results from pre-patterns of intracellular receptor pools. To address this question, living open-books where fixed, permeabilized and incubated with antibodies to reveal the total pool of pHLuo at high resolution in STED microscopy (**Fig. 6a,b,c**). Then, we quantified the receptor distribution within the growth cone. Interestingly, both PlxnA1 and Robo1 were observed to occupy similar growth cone areas and to have equal center of mass within the cone, thus excluding that polarized trafficking or storage pre-figure surface compartmentalization (**Fig. 6d**). Rather this supports that PlxnA1 and Robo1 are either delivered at the surface of different growth cone sub-domains or that their partitioning arises from membrane diffusion (**Fig.6e**).

**Fig. 6:**
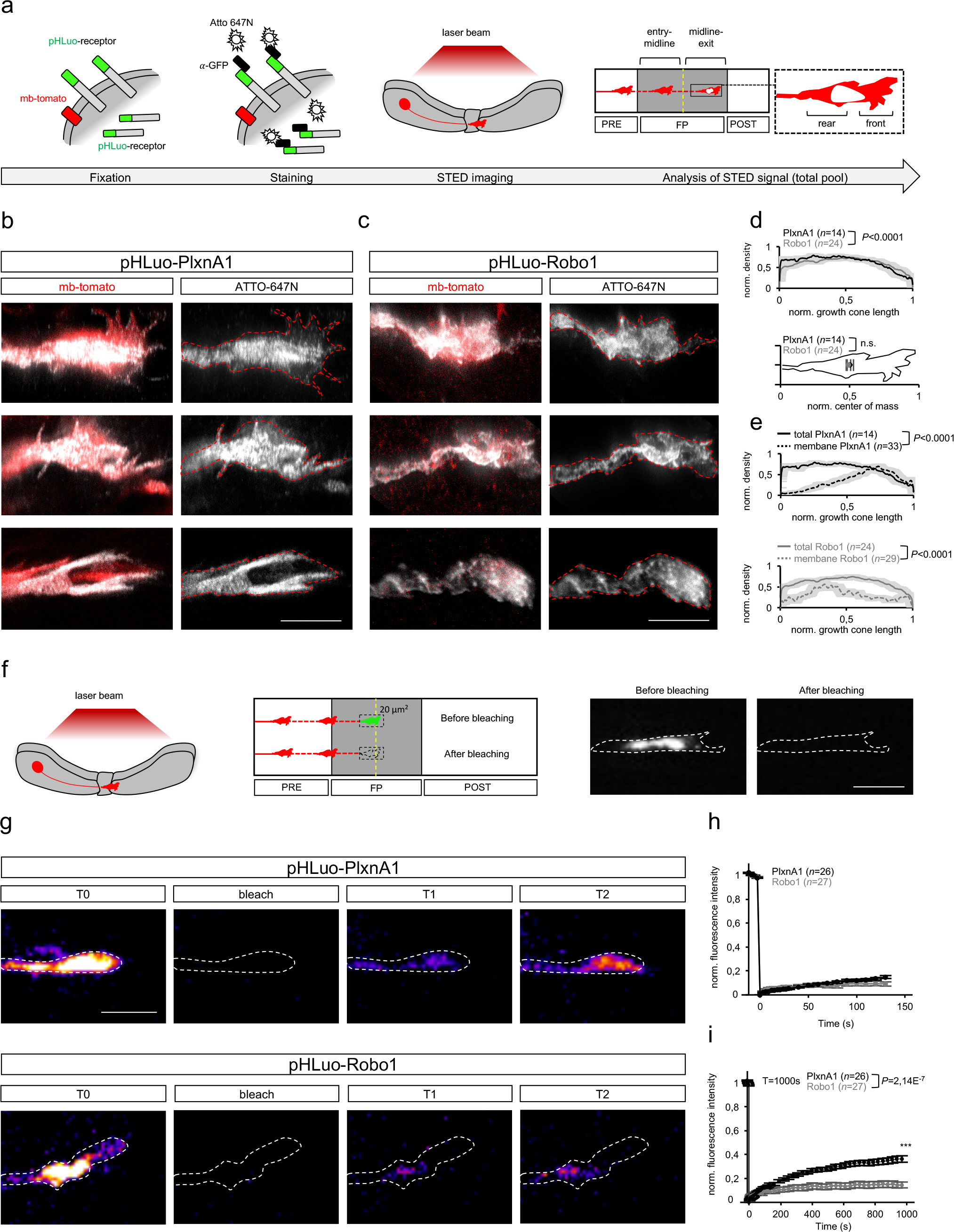
PlxnA1 and Robo1 partitioning at the cell surface is not generated *via* pre-patterns of intracellular receptor pools. Super resolution procedure for total receptor pool detection. Before STED imaging, open-books of embryos electroporated with pHLuo vectors were fixed, permeabilized and then immunolabeled with a primary anti-GFP antibody recognizing pHLuorin, followed by an ATTO-647N-conjugated secondary antibody. pHLuo density and distribution were analyzed in growth cones at different steps of FP navigation. The growth cone was segmented into front and rear sub-domains. (**b,c**) Microphotographs of representative commissural growth cones delineated with mb-tomato and labeled with ATTO-647N-conjugated GFP nanobodies. (**d**) Distribution of total pHLuo within the growth cones. Upper panel: Densities of total pHLuo-Plxna1 and -Robo1 signals normalized to the growth cone length. Lower panel: positions of pHLuo-PlxnA1 and -Robo1 center of mass normalized to growth cone lengths. Note that PlxnA1 and Robo1 total pool have similar center of mass within the growth cone (N PlxnA1=14 growth cones; N Robo1=24 growth cones. Error bars indicate mean ± SEM. Upper panel *P* is from Kolmogorov-Smirnov (KS) test; Lower panel *P* is from Mann-Whitney test). (**e**) Densities of membrane and total pHLuo in pHLuo-PlxnA1 (upper panel) and in pHLuo-Robo1 (lower panel) electroporated conditions, normalized to the growth cone length. Note the differential distribution along the growth cone rear-front axis between the two receptor pools (N membrane PlxnA1=33 growth cones; N total PlxnA1=14 growth cones; N membrane Robo1= 29 growth cones; N total Robo1=24 growth cones. *P* is from Kolmogorov-Smirnov (KS) test). (**f**) FRAP procedure. Growth cones navigating the FP and expressing either pHLuo-PlxnA1 or pHLuo-Robo1 were bleached using a 488nm diode laser and fluorescence recovery was monitored. The dashed rectangle indicates the typical bleached area. (**g**) Representative images from time-lapse sequences illustrating the bleaching and the fluorescence recovery of pHLuo-PlxnA1 and pHLuo-Robo1 in the growth cone. T1=370s, T2=970s. (**h,i**) Graphs of fluorescence recovery for pHLuo-PlxnA1 and pHLuo-Robo1. Dots represent the mean of three independent experiments (PlxnA1: N=26 growth cones; Robo1: N=27 growth cones. Error bars indicate mean ± SEM. *P* is from Student t test performed on values at 1000s). Scale bars in **b**, 5μm; in **f** and **g**, 10μm.

To get further insights into PlxnA1 and Robo1 sorting, we set-up a paradigm of FRAP experiments, targeting commissural growth cones expressing pHLuo-receptors and navigating the FP in living open-books. The pHLuo of receptor pool present in the intracellular acid compartment is protected from bleaching-induced damages and can thus produces fluorescence which then reports membrane insertion events (Ashby et al, 2004). To specifically assess the dynamics of membrane insertion of PlxnA1 and Robo1, the pHLuo-receptor fluorescence in an area of 15 to 20μm^2^ covering the entire growth cone surface was bleached at 80-90% (see methods) (**Fig. 6f,g**). As expected from this paradigm, at short 100 and 200 seconds time points, contribution of lateral membrane diffusion was negligible for both receptors, with a fluorescence recovery representing less than 0.5% of the initial fluorescence (**Fig. 6h**). Interestingly over a period of 17 minutes, we observed that Robo1 recovery was modest, rapidly reaching a plateau at around 17% of the initial fluorescence level. In contrast, the recovery level of PlxnA1 was significantly higher (**Fig. 6i, Supplementary Movies 11,12 for PlxnA1 and Supplementary Movies 13,14 for Robo1**). It did not reach yet a plateau and attained 38% of the initial fluorescence at the end of the recording period. Thus, the dynamics of membrane insertion strikingly differs between Robo1 and PlxnA1. While intracellular PlxnA1 can be rapidly mobilized for externalization, Robo1 intracellular pool might be almost depleted when commissural growth cones sort Robo1, strongly limiting additional supply of receptors at the surface.

### PlxnA1 but not Robo1 intracellular domain encodes receptor temporal patterns

Lastly, we investigated how could be dictated the specific timing of cell surface PlxnA1 and Robo1 sorting when commissural growth cones get exposed to local FP cues. We thought to construct chimeric receptors in which the extracellular (ECD) and intracellular domains (ICD) were swapped (**Fig. 7a**). pHLuo-PlxnA1^ECD^-Robo1^ICD^ and pHLuo-Robo1^ECD^-PlxnA1^ICD^ were introduced in the neural tube of chicken embryos and the position of growth cones that turned on the pHLuo fluorescence was plotted to build cartographies of receptor cell surface sorting. We found that the specific temporal profile of PlxnA1 sorting at the FP entry was preserved in context where the chimeric receptor contains PlxnA1 ICD but not ECD (**Fig. 7b,c,d,e**). In contrast for Robo1, neither Robo1 ICD nor ECD alone was sufficient to confer the temporal sorting of the native receptor in the second half of the FP navigation. Moreover, PlxnA1^ECD^-Robo1^ICD^ condition led to premature sorting of the receptor prior to FP entry, thus also revealing that PlxnA1 ICD is required for preventing the native receptor to be sorted at the pre-crossing stage (**Supplementary Movies 15,16 for pHluo-PlxnA1^ECD^-Robo1^ICD^ and Supplementary Movies 17,18 for pHluo-Robo1^ECD^-PlxnA11^ICD^**).

**Fig. 7:**
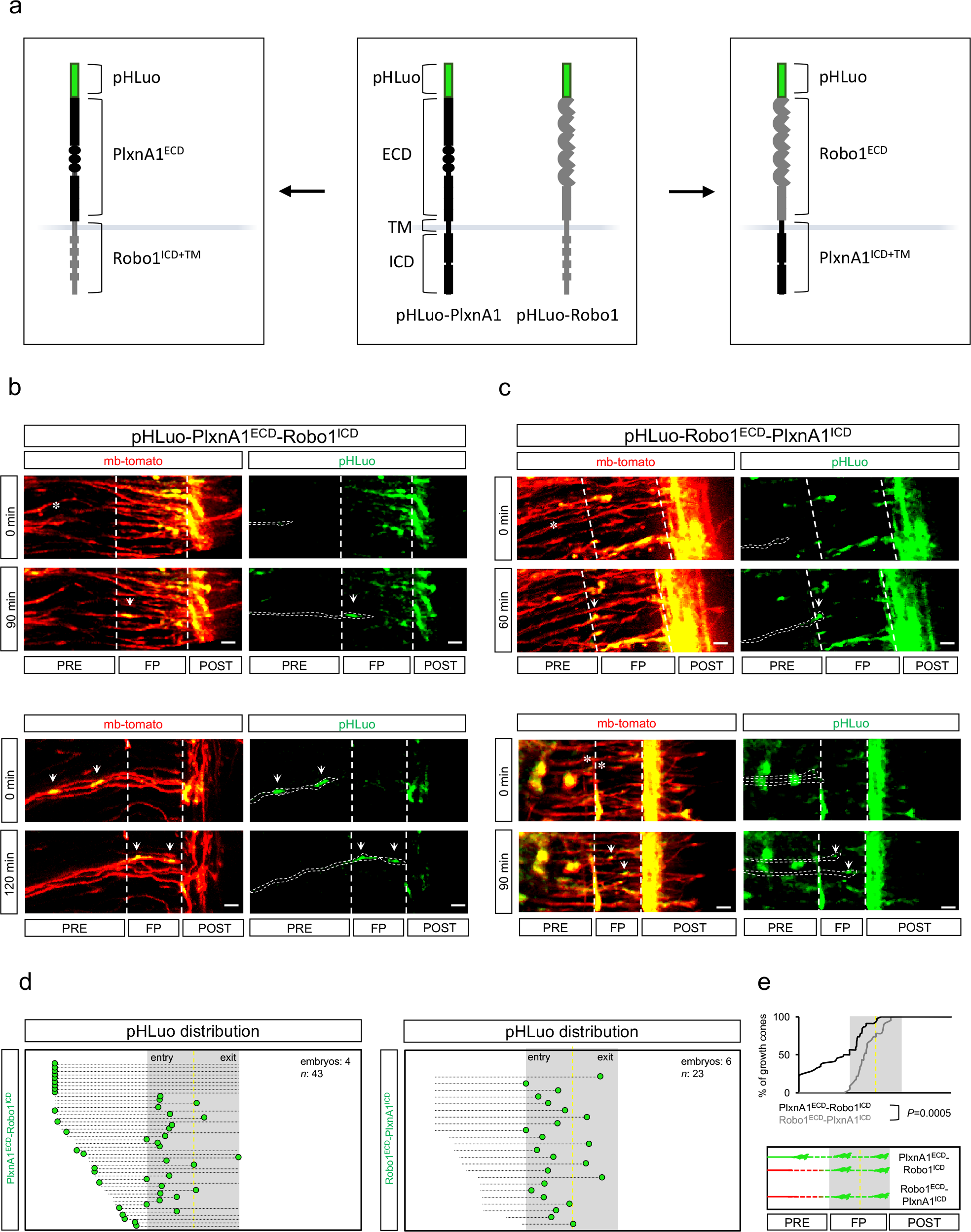
PlxnA1 but not Robo1 intracellular domain encodes membrane insertion temporal pattern. (**a**) Representation of pHLuo-PlxnA1 and pHluo-Robo1 receptor structures. The pHluo-PlxnA1-Robo1 chimera (left panel) consists of the PlxnA1^ECD^ fused to Robo1 transmembrane (TM) and ICD. The pHLuo-Robo1-PlxnA1 chimera consists of the Robo1^ECD^ fused to PlxnA1 TM and ICD. (**b**) Time-lapse sequences illustrating the dynamics of pHLuo-PlxnA1-Robo1 chimera. Arrowheads point pHLuo detected in growth cones at the FP entry (upper panel) and in growth cones prior to their arrival to the FP (lower panel). (**c**) Time-lapses sequences showing the dynamics of pHLuo-Robo1-PlxnA1 chimera. The flashes are detected at the FP entry. (**d**) Cartographies of pHLuo-PlxnA1^ECD^- Robo1^ICD^ (left panel) and pHLuo-Robo1^ECD^- PlxnA1^ICD^ (right panel) chimera dynamics from movie analysis. Dashed lines indicate the overall trajectory of single growth cones and green spots the first pHLuo detection. Note that for both chimeras, flashes are detected at the FP entry. However, for pHLuo-PlxnA1^ECD^-Robo1^ICD^ chimera, pHLuo is also detected in growth cones prior to their crossing. (**e**) Upper panel: cumulative fractions showing differential dynamics of pHLuo-PlxnA1^ECD^-Robo1^ICD^ and pHLuo-Robo1^ECD^-PlxnA1^ICD^ dynamics during FP navigation (*P* value is from the Kolmogorov-Smirnov (KS) test). Lower panel: Summary of the temporal sequence of PlxnA1^ECD^-Robo1^ICD^ and pHLuo-Robo1^ECD^-PlxnA1^ICD^. Scale bars in **b** and **c**, 10 μm.

## Discussion

During neural circuit wiring, focal and timed patterns of signaling are thought to be crucial for the specification of axonal trajectories but are yet highly challenging to visualize since this requires complex experimental paradigms preserving the topography of guidance molecules and their proper perception by the growth cones. Our live imaging and STED microscopy set-up thus pave the way to access the temporal and spatial resolution of signaling activities in neuronal growth cones facing guidance decisions in their physiological navigation environment. We report here that beyond their synthesis and trafficking to the axon, guidance receptors are exposed at the surface of the growth cones at precise timing and location. During commissural axon navigation, these receptor-specific cell surface codes can thus shape spatially and temporally distinct signaling from coincident midline cues, allowing their concerted action. Such a dynamic regulation of receptor equipment at the cell surface might be exploited in a variety of biological processes during which cells must adapt their perception of extracellular cues in a context-dependent manner. Particularly, accommodating fast and flexible perception of extracellular signals is indeed a prerequisite for cells which, as the axons do so, migrate along highly stereotyped and long-distance pathways, getting exposed to fluctuating combinations of guidance cues (Te Boekhorst *et al.*, 2016).

In the context of commissural axon navigation, our results bring new insights and directions for further investigation of the functional outcome resulting from the different receptor-mediated signaling. Our data report a temporal sequence of Nrp2, PlxnA1, Robo1 and Robo2 surface sorting, which equip commissural growth cones at successive steps of their navigation. PlxnA1/Nrp2-mediated Sema3B and PlxnA1-mediated SlitC activities can thus be switched on from the FP entry, while Robo1-mediated SlitN signaling might start after the midline, and Robo2-mediated signaling at a next post-crossing choice point. Our analysis from super resolution microscopy showing that Robo1 and PlxnA1 receptors have distinct distribution at the growth cone surface gives complementary information on the organization of the repulsive activities mediating FP crossing. It could be that Slits and Sema3B repellents have synergistic contributions to growth cone progression across the FP, receptor compartmentalization concentrating their signaling onto different sub-domains of the growth cone to promote its forwards progression. Nevertheless, and also supported by the distinct temporal patterns of their receptors, the repellent activities might rather be uncoupled, regulating different steps or aspects of the growth cone navigation through distinct downstream signaling cascades. Our live imaging analysis of growth cone behavior prior to and after receptor sorting also supports that signaling via these receptors have different functional outcome. Interestingly, the gain of Robo1 coincided with increased rostro-caudal exploration behavior of the growth cones. This did not result in premature deviation of the trajectory in the FP, but appeared rather to pre-figure the longitudinal turning decision that will be taken at the FP exit. In contrast, modification of exploratory potential was never observed after gain of membrane PlxnA1, despite the evidence from over-expression conditions that the receptor is functionally active. This supports that signaling by PlxnA1 and Robo1 ligands have different outcome. The manifestation of PlxnA1-mediated signaling on the growth cone behavior remains to be uncovered. Whatever the possibilities, its outcome might be to prevent the growth cone from turning back since genetic deletion of PlxnA1 in mouse embryos results in re-crossing phenotypes (Delloye-Bourgeois *et al.*, 2015).

The front-rear partitioning of PlxnA1 and Robo1 receptors also provides some clues into the mode of action of their respective ligands. Such compartmentalization correlates with the spatial organization of growth cone cytoskeletal components and could serve as a mechanism for generating focal signaling onto different cytoskeletal components. Indeed, growth cone responses to guidance cues rely on regulations of the dynamics of both microtubule and actin cytoskeleton, which physically mainly occupy the central and the filopodia-rich peripheral domains, respectively (Cammarata *et al*., 2016; Kahn and Baas, 2016).

Finally, our findings that the membrane sorting of Robo1 and Robo2 coincides with changes of growth cone behaviors, that are activated at different time points of their navigation, strengthen the view that temporally controlling the availability of the receptors at the growth cone surface is a general mechanism to accommodate the growth cones to novel guidance challenges.

These findings also highlight unexpected differences between Robo receptors. Despite evidence in several systems that Robo1 and Robo2 have distinct contributions (Kim *et al*., 2011), which specific guidance decisions they control during commissural axon navigation is yet unclear. Both Robo1 and 2 receptors transduce Slit signals and are expressed by commissural neurons. Both have been reported expressed at low levels in pre-crossing/crossing commissural axons and enriched at the post-crossing stage (Long *et al*., 2004).

In the mouse, Robo1 but not Robo2 deletion alters FP crossing. Nevertheless, double Robo deletion was reported to aggravate the impact of Robo1 invalidation (Jaworski *et al*., 2010). In contrast, specifically in Robo2^−/−^embryos, commissural axons failed to reach the lateral funiculus, consistent with reported Robo2 enrichment in post-crossing lateral axon segments (Long *et al*., 2004). Dominant-negative based approach to abrogate Robo signaling in the chick embryo also resulted in alteration of the post-crossing lateral navigation (Reeber *et al*., 2008). In the drosophila ventral cord, post-crossing commissural tracts segregate at distinct lateral positions from the midline, as a result of tract-specific combinations of Robo receptors conferring them different sensitivity to midline Slits (Neuhaus-Follini and Bashaw, 2015). As a parallel, combinations of Robo receptors setting distinct responses to FP Slits are thought to control the position of post-crossing tracts navigating within the ventral and lateral funiculi in the mouse (Long *et al*., 2004; Jaworski *et al*., 2010). Our study enlightens drastic differences between Robo1 and Robo2 receptor sorting that could not be anticipated from their general expression patterns. In contrast to Robo1, Robo2 is sorted only in growth cones navigating the LF and accomplishing their longitudinal turning. Such temporally accurate Robo2 sorting is unlikely involved in the perception of FP Slits. It rather suggests that the growth cone acquire perception of a guidance cue precisely at the VF/LF transition, which calls to question the exact mechanisms controlling medio-lateral position choices of post-crossing axons.

Overall, we uncovered at different choice points strong and specific regulations of membrane sorting for three major guidance receptors known to contribute to the pathfinding of multiple neuronal tracts (Jongbloets and Pasterkamp, 2014; Blockus and Chédotal, 2016). Thus, dynamic control of cell surface expression might have general and crucial contribution to axon navigation. Mechanisms controlling the cell surface addressing of PlxnA1 and Robo1 in vertebrates have been proposed, implicating regulated exocytosis and protein processing (Nawabi *et al*., 2010; Charoy *et al*., 2012; Philipp *et al*., 2012). Our findings add another piece of information, showing that Robo1 sorting depends on local signals released by the FP. The identity of these signals remains to be determined. Whether they are receptor-specific or similar to those that trigger PlxnA1 and Robo3.2 is yet unknown. Whatever the case, our study demonstrates that the temporal differences of Robo1 and PlxnA1 are dictated by receptor-specific membrane insertion dynamics. First the temporality of both Robo1 and PlxnA1 surface delivery is controlled at the step of membrane addressing. Indeed, in both cases, we observed receptors present in the growth cones but not yet sorted at their surface. Second, in our FRAP experiments, depleting pHLuo fluorescence in growth cones navigating the FP in open-books and comparing the timing of fluorescence recovery revealed striking differences between Robo1 and PlxnA1 dynamics. This might reflect differences in protein availability in the growth cone, which could be set through translational and post-translational protein regulations as well as through distinct protein trafficking pathways. Robo1 sorting from intracellular vesicles was reported to occur through up-regulation of RabGDI (Philipp *et al*., 2015). It would be interesting to determine whether Robo1 temporal pattern is conditioned by the assembly of functional membrane delivery machinery. Control of Robo1 receptor pool in commissural neurons was also recently reported to occur via microRNA-mediated regulation of Robo1 synthesis (Yang *et al.*, 2018). This regulation is unlikely responsible for specifying the timing of surface sorting since our expression vectors do not encode the regulatory sequences underlying microRNA regulation. We previously reported that regulation of PlxnA1 sorting depends on processing by calpains, with integral but not cleaved PlxnA1 undergoing cell surface sorting (Nawabi *et al.*, 2010; Charoy *et al*., 2012). Processing suppression resulting from calpain inhibition, which we found to occur when commissural axons get exposed to local FP signals, would allow fast membrane insertion of integral PlxnA1, which is consistent with our findings from FRAP experiments.

Our analysis of the total and membrane receptor pools in STED microscopy also suggests that the sorting mechanism not only controls the timing of membrane availability of the receptors but also their spatial distribution at the growth cone surface. Such membrane compartmentalization could emerge from polarized receptor delivery or rather through rearrangements at the membrane through selective retention or retrieval.

Overall, our study suggests that the generation of specific temporal sequences of receptor cell surface sorting represent a mechanism whereby commissural growth cones discriminate simultaneous signals functionalizing them at precise timing during spinal cord navigation.

## Supporting information

Supplementary Movie1 Nrp2. Related to Figure 1avi

Supplementary Movie2 Nrp2. Related to Figure 1

Supplementary Movie3 PlexA1.Related to Figure 1

Supplementary Movie4 PlexA1.Related to Figure 1

Supplementary Movie5 Robo1.Related to Figure 1

Supplementary Movie6 Robo1.Related to Figure 1

Supplementary Movie7 PlexA1 over expression 4ug ul. Related to FigureS 1

Supplementary Movie8 Robo1 over expression 4ug ul. Related to Figure S1

Supplementary Movie9 Robo2. Related to Figure 3

Supplementary Movie10 Robo2.Related to Figure 3

Supplementary Movie11 FRAP pHLuo-PlxnA1. Related to Figure 6

Supplementary Movie12 FRAP pHLuo-PlxnA1. Related to Figure 6

Supplementary Movie13 FRAP pHLuo-Robo1. Related to Figure 6

Supplementary Movie14 FRAP pHLuo-Robo1. Related to Figure 6

Supplementary Movie15 ECDPlxnA1-ICDRobo1. Related to Figure7

Supplementary Movie16 ECDPlxnA1-ICDRobo1. Related to Figure 7

Supplementary Movie17 ECDRobo1-ICDPlxnA1. Related to Figure 7

Supplementary Movie18 ECDRobo1-ICDPlxnA1. Related to Figure 7

## Acknowledgments

We thank A. Chedotal for sharing Robo mouse model, E. Derrington for input on the manuscript, O. Raineteau for scientific discussions. STED microscopy was done in the Bordeaux Imaging Center (BIC), CNRS-INSERM-Bordeaux University, member of the national infrastructure France BioImaging supported by the French National Research Agency (ANR-10-INBS-04). We thank Christel Poujol and Magali Mondin (BIC) of Bordeaux for advices. This work was conducted within the frame of the LabEx CORTEX and DEVWECAN of Université de Lyon, within the program “Investissements d’Avenir” (ANR-11-IDEX-0007) operated by the French National Research Agency (ANR). The study was supported by an ANR funding to VC and OT, and the Association Française contre les Myopathies (AFM) to VC.

## Author contributions

conceptualization: VC; methodology: AP, HD, JF, LB, VC, OT; investigation AP (live imaging and STED microscopy), HD (STED microscopy and FRAP experiments), KK and MB (open-book assays, molecular biology, tool validation), STD and TG (chimeric receptor constructions); writing AP, VC; writing editing: HD, JF; formal analysis: AP, HD, JF, OT, VC.

## Competing interests

Authors declare no competing interests.

## References

Ashby, M.C., De La Rue, S.A., Ralph, G.S., Uney, J., Collingridge, G.L., Henley, J.M. (2004). Removal of AMPA receptors (AMPARs) from synapses is preceded by transient endocytosis of extrasynaptic AMPARs. J Neurosci 24, 5172–6.

Blockus, H., Chédotal, A. (2016). Slit-Robo signaling. Development 143, 3037–44.

Cammarata, G.M., Bearce, E.A., Lowey, L.A. (2016). Cytoskeletal social networking in the growth cone: How +TIPs mediate microtubule-actin cross-linking to drive axon outgrowth and guidance. Cytoskeleton (Hoboken) 74, 461–76.

Charoy, C., Nawabi, H., Reynaud, F., Derrington, E., Bozon, M., Wright, K., Falk, J., Helmbacher, F., Kindbeiter, K., Castellani, V. (2012). GDNF activates midline repulsion by Semaphorin3B via NCAM during commissural axon guidance. Neuron 75, 1051–66.

Chen, Z., Gore, B.B., Long, H., Ma, L., Tessier-Lavigne, M. (2008). Alternative splicing of the Robo3 axon guidance receptor governs the midline switch from attraction to repulsion. Neuron 58, 325–332.

Colak D., Ji S.J., Porse B.T., Jaffrey S.R. (2013). Regulation of Axon Guidance by Compartmentalized Nonsense-Mediated mRNA Decay. Cell 153, 1252–1265.

Delloye-Bourgeois, C., Jacquier, A., Charoy, C., Reynaud, F., Nawabi, H., Thoinet, K., Kindbeiter, K., Yoshida, Y., Zagar, Y., Kong, Y., Jones, Y.E., Falk, J., Chédotal, A., Castellani, V. (2015). PlexinA1 is a new Slit receptor and mediates axon guidance function of Slit C-terminal fragments. Nat Neurosci 18, 36–45.

Delloye-Bourgeois, C., Jacquier, A., Falk, J., Castellani, V. (2014). Use of pHLuorin to assess the dynamics of axon guidance receptors in cell culture and in the chick embryo. J Vis Exp (83):e50883. doi: 10.3791/50883.

Evans, T.A., Bashaw, G.J. (2010). Axon guidance at the midline: of mice and flies. Curr Opin Neurobiol. 20, 79–85.

Jacob, T.C., Bogdanov Y.D., Magnus C., Saliba R.S., Kittler J.T., Haydon P.G., Moss S.J. (2005). Gephyrin regulates the cell surface dynamics of synaptic GABAA receptors. J Neuroscience 25, 10469–78.

Jaworski, A., Long, H., Tessier-Lavigne, M. (2010). Collaborative and specialized functions of Robo1 and Robo2 in spinal commissural axon guidance. J Neurosci 30, 9445–9453.

Jongbloets, B.C., Pasterkamp, R.J. (2014). Semaphorin signaling during development. Development 141, 3292–7.

Kahn, O.I., Baas, P.W. (2016). Microtubules and Growth Cones: Motors Drive the Turn. Trends Neurosci 39, 433–440.

Kim, M., Roesener, A.P., Mendonca, P.R., Mastick, G.S. (2011). Robo1 and Robo2 have distinct role in pioneer longitudinal axon guidance. Dev Biol. 358, 181–8.

Long, H., Sabatier, C., Ma, L., Plump, A., Yuan, W., Ornitz, D.M., Tamada, A., Murakami, F., Goodman, C.S., Tessier-Lavigne, M. (2004). Conserved roles for Slit and Robo proteins in midline commissural axon guidance. Neuron 42, 213–223.

Nawabi, H., Briancon-Marjollet, A., Clark, C., Sanyas, I., Takamatsu, H., Okuno, T., Kumanogoh, A., Bozon, M., Takeshima, K., Yoshida, Y., Moret, F., Abouzid, K., Castellani, V. (2010). A midline switch of receptor processing regulates commissural axon guidance in vertebrates. Genes Dev. 24, 396–410.

Neuhaus-Follini., A. Bashaw. G.J. (2015). Crossing the embryonic midline: molecular mechanisms regulating axon responsiveness at an intermediate target. Wiley Interdiscip Rev Dev Biol. 4(4):377–89

Philipp, M., Niederkofler, V., Debrunner, M., Alther, T., Kunz, B., Stoekli, E.T. (2012). RabGDI controls axonal midline crossing by regulating Robo1 surface expression. Neural Dev. 9, 7–36.

Pignata, A., Ducuing, H., Castellani, V. (2016). Commissural axon navigation: Control of midline crossing in the vertebrate spinal cord by the semaphorin 3B signaling. Cell Adh Migr. 10, 604–617.

Reeber, S.L., Sakai, N., Nakada, Y., Dumas, J., Dobrenis, K., Johnson, J.E., Kaprielian, Z. (2008). Manipulating Robo expression *in vivo* perturbs commissural axon pathfinding in the chick spinal cord. J Neurosci. 28, 8698–8708.

Stoekli, E.T. (2018). Understanding axon guidance: are we nearly there yet? Development 145(10).

Te Boekhorst V., Preziosi L., Friedl P. (2016). Plasticity of Cell Migration In Vivo and In Silico. Annu Rev Cell Dev Biol. 6, 491–526.

Wilson S.I., Shafer B., Lee K.J., Dodd J. (2008). A molecular program for contralateral trajectory: Rig-1 control by LIM homeodomain transcription factors. Neuron. 59, 413–24.

Yang T., Huang H., Shao Q., Yee S., Majumder T., Liu G. (2018). miR-92 Suppresses Robo1 Translation to Modulate Slit Sensitivity in Commissural Axon Guidance. Cell Reports 24, 2694–2708.

Zou, Y., Stoeckli, E., Chen, H., Tessier-Lavigne, M. (2000): Squeezing axons out of the gray matter: A role for slit and semaphorin proteins from midline and ventral spinal cord. Cell 102, 363–375.

