## Supplementary Materials for "Spatial and temporal profiling of receptor membrane insertion controls commissural axon responses to midline repellents"

#### Supplementary Figure 1

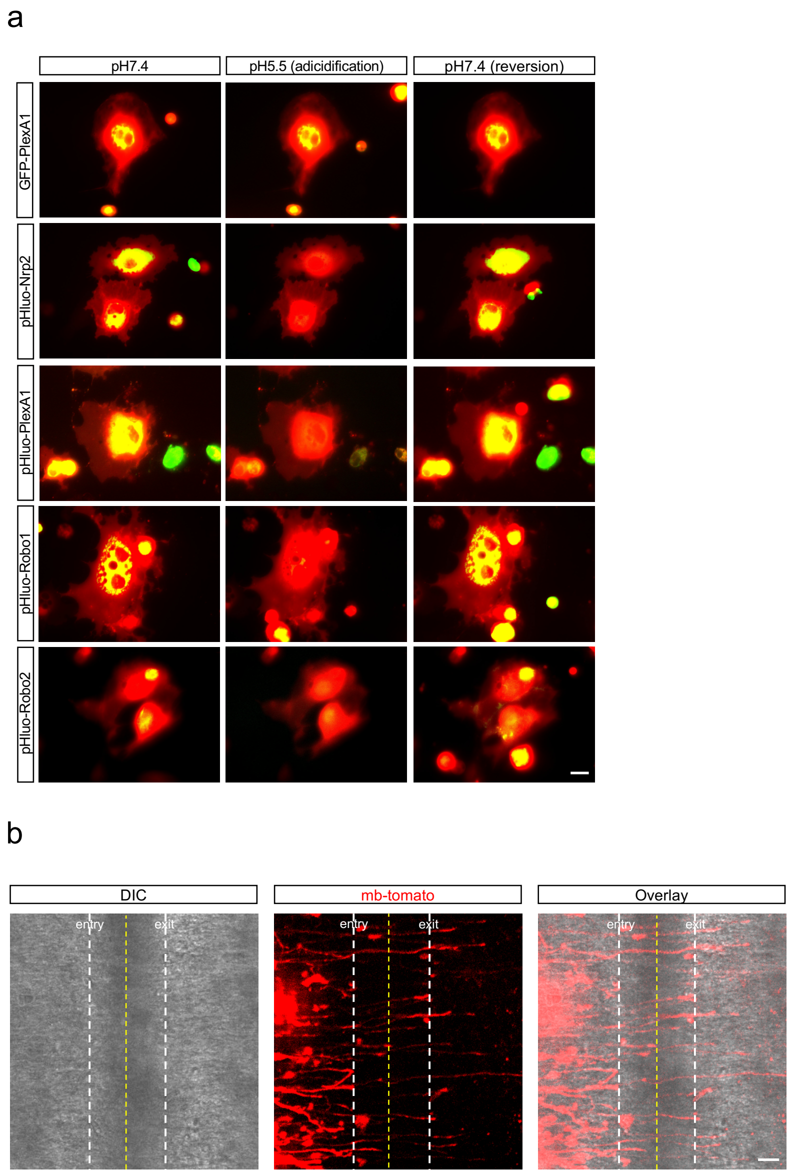


**Supplementary Figure 1: pH-dependency of receptor fluorescence and FP delimitation in open-books**

(**a**) Cos7 cells transfected with pHLuo receptors and live monitored using confocal microscopy. pHLuo receptors fluoresce at the membrane when cells are cultivated in a pH7.4 culture medium. When the culture medium was acidified up to pH5.5, pHLuo lose its fluorescence. pH reversion to neutrality restored the cell surface fluorescence. These pH-dependent properties were not observed in control GFP-PlexA1 transfected COS cells. Scale bars, 5 µm. (**b**) Spinal cord morphology was revealed using DIC to delimit FP entry and exit. Scale bars, 50 µm.

### Supplementary Figure 2

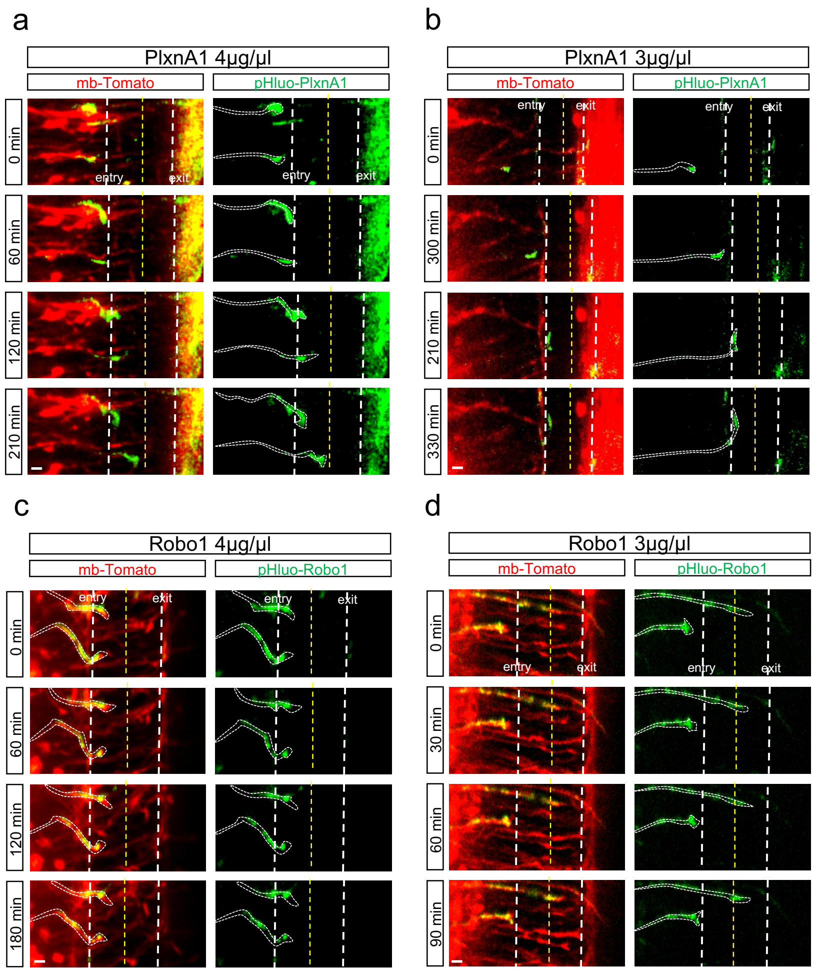


**Supplementary Figure 2**: **Over-loading commissural growth cones with receptors alters their FP navigation**

(**a-d**) Time-lapse sequences showing over-expression phenotypes induced by the electroporation of high doses of pHLuo-PlxnA1 or pHLuo-Robo1 plasmids (4µg/µl and 3µg/µl). Harrow-heads point growth cones sorting prematurely the receptors, which stall or aberrantly turn at the FP entry. Scale bars in **a-d**, 10 µm.

**Supplementary Movies 1-2: Dynamics of pHLuo-Nrp2**

pHLuo-Nrp2 is exposed at the commissural growth cone surface from the pre-crossing stage and remains over entire FP crossing. White arrows point the growth cones during FP navigation. FP: floor plate.

**Supplementary Movies 3-4: Dynamics of pHLuo-PlexA1**

pHLuo-PlxnA1 is addressed to the surface when commissural growth cones navigate the first half of the FP, from the FP entry point to the midline. White arrows point the growth cones during FP navigation. FP: floor plate.

**Supplementary Movies 5-6: Dynamics of pHLuo-Robo1**

pHLuo-Robo1 is sorted to the cell surface when commissural growth cones navigate the second half of the FP, from the midline to the FP exit point. White arrows point the growth cones during FP navigation. FP: floor plate.

**Supplementary Movie 7: Over-loading commissural growth cones with pHLuo-PlexA1 alters their FP navigation.**

Time-lapse sequences showing over-expression phenotypes induced by the electroporation of high doses of pHLuo-PlxnA1 plasmids (4µg/µl). White arrows point growth cones prematurely stalling at FP entry. FP: floor plate.

**Supplementary Movie 8: Over-loading commissural growth cones with pHLuo-Robo1 alters their FP navigation.**

Time-lapse sequences showing over-expression phenotypes induced by the electroporation of high doses of pHLuo-Robo1 plasmids (4µg/µl). White arrows point commissural growth cones prematurely exposing pHLuo-Robo1 and stalling at the FP entry. FP: floor plate.

**Supplementary Movies 9-10: Dynamics of pHLuo-Robo2**

pHLuo-Robo2 is sorted to the cell surface in post-crossing growth cones that chose to navigate in the lateral funiculus, turning longitudinally to the FP (LF). White arrows point the growth cones during the navigation in post-crossing compartment. FP: floor plate; VF: ventral funiculus; LF: lateral funiculus.

**Supplementary Movies 11-12: FRAP sequences of pHLuo-PlxnA1$^{+}$ growth cones.**

The pHLuo-receptor fluorescence in an area of 15 to 20µm$^2$ covering the entire growth cone surface was bleached at 80-90%. The recovery was measured over a period of 17 minutes.

**Movies S13-S14: FRAP sequences of pHLuo-Robo$^{+}$ growth cones.**

The pHLuo-receptor fluorescence in an area of 15 to 20µm$^2$ covering the entire growth cone surface was bleached at 80-90%. The recovery was measured over a period of 17 minute

**Movies S15: pHluo-PlxnA1$^{ECD}$-Robo1$^{ICD}$ chimera dynamics.**

pHluo-PlxnA1$^{ECD}$-Robo1$^{ICD}$ is exposed at the commissural growth cone surface since the pre-crossing stages. Flashes are also detected at the FP entry.

**Movies S17-S18: pHluoRobo1$^{ECD}$-PlxnA1$^{ICD}$ chimera dynamics**

pHluo-PlxnA1$^{ECD}$-Robo1$^{ICD}$ does not confer the temporal sorting of the native receptor in the second half of the FP navigation.

#### Materials and Methods

##### Receptor molecular biology

FL mouse pHLuo-PlxnA1 was generated by introducing in Nter the coding sequence of the pHLuo cloned from a vector encoding GABA A pHLuo-GFP (Jacob et al., 2005). pHLuo derived from this vector was fused to FL rattus Robo1 and Robo2 sequences, kindly provided by A. Chedotal laboratory, to obtain pHLuo-Robo1 and pHLuo-Robo2 vectors. Using the same strategy, FL mouse Nrp2 kindly provided by Püschel laboratory, was fused to pHLuo to obtain pHLuo-Nrp2 vectors. pHLuo-receptors where finally cloned into a PCAGEN vectors with an ires-mb-tomato sequence.

##### *In vitro* test of pH fluorescence dependency

Cos7 cells were plated in a glass-bottom dish 35mm in 2ml of complete Dulbecco's modified eagle medium (DMEM – 10% fetal bovine serum – 1 mM sodium pyruvate – 25 U/ml penicillin/streptomycin – 2,5 µg/ml Amphotericin B – pH7.4). 24h after, cells were transfected with 2 µg of plasmid encoding pHLuorin-tagged receptors and the transfection reagent was added. 48h later, live imaging of the cells was performed at 40X magnification. Cells were first imaged at pH 7.4 then, 1,25ml of pH3.5 complete DMEM was added to achieve a pH of 5.5 in the culture medium. Next, 1.2 of pH complete DMEM was injected to revert the pH of the medium to neutrality. Images were taken every 20 seconds for 10 minutes.

**_In ovo_ electroporation**

*In ovo* electroporation of HH14/HH15 chick embryos was performed as described previously (Delloye-Bourgeois *et al.,* 2014). Plasmids were diluted at the following concentration: 1.5 µg/µl pHLuo-Robo1-*ires*-tomato; 2.5 µg/µl pHLuo-Robo2-*ires*-tomato; 1 µg/µl pHLuo-Np2-*ires*-tomato; 2 µg/µl pHLuo-PlexA1 and 0.3 µg/µl mb-tomato. Plasmids were diluted in UP $H_2O$ and the solution was injected into the lumen of the neural tube using picopritzer III (Micro Control Instrument Ltd., UK). Electrodes (CUY611P7-4, Sonidel) were placed along the back of the embryo, at the thoracic level, and 3 pulses (25V, 500ms interpulse) were delivered by CUY-21 generator (Sonidell). Electroporated embryos were then incubated at 38.5°C.

**Open book culture**

48 hours after electroporation, embryos at HH25/HH26 were harvested in cold HBSS and the spinal cords were dissected. Spinal cords were mounted in 0.5% agarose diluted in F12 medium and placed on glass bottom dishes (P35G-1.5-14-C, MatTek). After agarose solidification, spinal cords were overlaid with 3ml of F12 medium supplemented with 10% FCS (F7524; Sigma-Aldrich), 1% Penicillin/Streptomycin (Sigma-Aldrich) and 20mM HEPES buffer (15630-049, ThermoFischer Scientific).

**Mouse spinal cord electroporation and culture**

E12 mice embryos were collected and fixed on a SYLGARD (Dow Corning) culture plate in Leibovitzs 15 medium (ThermoFisher) supplemented with Glucose 1M (Sigma-Aldrich). Plasmids were injected into the lumen of the neural tube using picopritzer III (Micro Control Instrument Ltd., UK). Electrodes (CUY611P7-4, Sonidel) were placed along the back of the embryo, at the thoracic level, and 3 pulses (25V, 500ms interpulse) were delivered by CUY-21 generator (Sonidell). Spinal cords were dissected from the embryos and cultured on Nucleopore Track-Etch membrane (Whatman) for 48 hours in Slice Culture Medium (*Polleux and Ghosh., 2002*).

**Live imaging and data analysis**

Live imaging was performed with an Olympus IX81 microscope equipped with a spinning disk

(CSU-X1 5000 rpm, Yokogawa) and Okolab environmental chamber maintained at 37°C. Image were acquired with a 20X objective by EMCCD camera (iXon3 DU-885, Andor technology). Usually for spinal cord culture, 15-30 planes spaced of 0,5-3µm were imaged for each spinal cord at 30-minute interval for 10 hours approximatively. To reduce exposure time and laser intensity, acquisitions were done using binning 2x2. Images were acquired using IQ3 software using multi-position and Z stack protocols. Z stack projections of the movies were analyzed in ImageJ software. The analysis of pHLuo-flashes was performed from time-lapse acquisitions *in vivo*. For cartography representation, the lengths of PRE-crossing and POST-crossing compartment were normalized on FP length.

**Detection of the total pool of pHLuorin**

48h after electroporation, at HH25/HH26, the embryos were harvested in cold HBSS and the spinal cords were dissected and fixed for 2 hours with PBS supplemented with 4% paraformaldehyde (PFA). The % of the commissural population expressing total pool of pHLuo in PRE-crossing was calculated by qualitative analysis of Z stack projections. The length of the PRE-crossing segments expressing total pHLuo was measured with Image J software.

**Explant cultures**

FPs were isolated from HH25/HH26 chick embryos and cultured in tridimensional plasma clots in B27-supplmented Neurobasal medium (GIBCO). The supernatant ($FP^{cm}$) was collected after 48h. Electroporated spinal cord were dissected, cut into explants and left retrieve for 30min at 37°C. Then explants were placed on glass bottom dishes, previously coated with 10µg/ml Laminin and 50µg/ml polylysine and cultured for approximately 30h at 37°C in F12 medium supplemented with 0.4 Methylcellulose, 1X B27, 100ng/ml Netrin, 1/1000 Penistreptomycin. Explants were imaged at T0, then $FP^{cm}$ or Ctrl medium were used for the treatment and 20min after a second time point was recorded.

**Atto647N staining for surface receptor pool**

Spinal cords were incubated at 38°C for 20 minutes with F12 medium supplemented with 5% FCS (F7524; Sigma-Aldrich), 20mM HEPES buffer (15630-049, ThermoFischer Scientific) and 1/100 GFP-nanobodies Atto647N. Spinal cords were then rinsed 4 times with the same medium (not containing the GFP-nanobodies) and were fixed at room temperature for 2 hours with PBS supplemented with 4% paraformaldehyde (PFA) and 1% BSA (A7638 Sigma-Aldrich).

**Atto647N staining for total receptor pool**

Spinal cords were fixed at room temperature for 2 hours with PBS supplemented with 4% paraformaldehyde (PFA) and 6% BSA (A7638 Sigma-Aldrich). Spinal cords were then incubated 18h at room temperature with 1% BSA, rabbit 1/400 anti-GFP antibody (Invitrogen) and incubated over night at 4°C with 3% BSA, anti-rabbit ATTO647N.

**STED imaging and data analysis**

The staining was observed with a STED microscope (TCS SP8, Leica). STED illumination of ATTO 647N was performed using a 633-nm pulsed laser providing excitation, and a pulsed bi-photon laser (Mai Tai; Spectra-Physics) turned to 765 nm and going through a 100-m optical fiber to enlarge pulse width (100ps) used for depletion. A doughnut-shaped laser beam was achieved through two lambda plates. Fluorescence light between 650 and 740 nm was collected using a photomultiplier, using a HCX PL-APO CS 100/1.40 NA oil objective and a pinhole open to one time the Airy disk (60mm). Images were acquired with using Leica microsystem software and a Z stack protocol. Usually 10-20 planes spaced of 0,5µm where imaged for each growth cone. The growth cone perimeter was outlined basing on the mb-tomato signal. Average density of pHLuo-receptors in the growth cones was calculated from Z stack projections with Matlab software. Particle numbers and surfaces were calculated from Z stack projections with image J.

**Fluorescence Recovery After Photobleaching**

FRAP experiments were performed on spinal cord open-books electroporated with either pHLuo-Robo1-ires-mb-tomato or pHLuo-PlxnA1 and mb-tomato using a Leica DMI6000 (Leica Microsystems, Wetzlar, Germany) equipped with a confocal Scanner Unit CSU-X1 (Yokogawa Electric Corporation, Tokyo, Japan) and a scanner FRAP system, ILAS (Roper Scientific, Evry, France). Images were acquired in both green and red channels using a 63X objective and an Evolve EMCCD camera (Photometrics, Tucson, USA). Growth cones located in the FP were first monitored for 9s each 3s and then bleached using a 488nm diode laser at full power. This resulted in an 80-90% loss of the signal at t=0. Fluorescence recovery was then monitored for 1030s with acquisitions every 3s for 30s, then every 10s for 100s, and finally every 30s for 900s. The images were corrected for background noise, residual fluorescence right after the bleach was set to zero, and recovery curves were normalized to the fluorescence lost after the bleach. No other corrections were applied since unbleached growth cone fluorescence showed no significant decay during the acquisition period.

**Chimeric receptor molecular biology**

The pHLuo-PlxnA1$^{ECD}$ and Robo1$^{ICD}$ fragments were amplified by PCR from pHLuo-PlxnA1 vector (forward pHLuo-PlxnA1$^{ECD}$ 5’-CATCATTTTGGCAAAGAATTCATGGGCTGGTTCACTGGGA-3’; reverse pHLuo-PlxnA1$^{ECD}$ 5’-CAATGAAGGCCGGCAGTGTCAGCAGGCT-3’) and pHLuo-Robo1-*ires*-tomato vector (forward Robo1$^{ICD}$ 5’-GACACTGCCGGCCTTCATTGCGGGCATC-3’; reverse Robo1$^{ICD}$ 5’-CGCGATATCCTCGAGGAATTTTAGCTTTCAGTTTCCTCTAATTC-3’), respectively. The pHLuo-Robo1$^{ECD}$ and PlxnA1$^{ICD}$ were amplified by PCR from pHLuo-Robo1-*ires*-tomato vector (forward pHLuo-Robo1$^{ECD}$ 5’-CATCATTTTGGCAAAGAATTCATGGGCTGGTTCACTGGGA-3’; reverse pHLuo-Robo1$^{ECD}$ 5’- CCACAATGGCTGGCTGCTTCACCACGTC-3’) and pHLuo-PlxnA1 (forward PlxnA1$^{ICD}$ 5’-GAAGCAGCCAGCCATTGTGGGTATCGGTGGT-3’; reverse PlxnA1$^{ICD}$ 5’- CGCGATATCCTCGAGGAATTTCAGCTGCTCAGGGCCAT-3’), respectively.

For both PlxnA1 and Robo1, the transmembrane domain was included in the ICD.

The amplified PCR fragments were cloned into the EcoRI site of pCAGEN (Addgene plasmid # 11160) using the In-Fusion HD Cloning Plus kit (Clontech, Mountain View, CA, USA) and following the manufacturer’s instructions.

**Statistics**

All embryos which normally developed and expressing pHLuo-vectors at the thoracic level were included in the analysis. Sample size and statistical significances are represented in each figure

and figure legend. For each set of data, normality was tested and Student t or Mann-Whitney tests were performed when the distribution was normal or not, respectively. Statistical tests were performed using Biosta-TGV (CNRS) and Prism 6 software.
